# Profiling of the most reliable mutations from sequenced SARS-CoV-2 genomes scattered in Uzbekistan

**DOI:** 10.1101/2022.02.10.479714

**Authors:** Mirzakamol S. Ayubov, Zabardast T. Buriev, Mukhammadjon K. Mirzakhmedov, Abdurakhmon N. Yusupov, Dilshod E. Usmanov, Shukhrat E. Shermatov, Khurshida A. Ubaydullaeva, Ibrokhim Y. Abdurakhmonov

## Abstract

Due to rapid mutations in the coronavirus genome over time and re-emergence of multiple novel variants of concerns (VOC), there is a continuous need for a periodic genome sequencing of SARS-CoV-2 genotypes of particular region. This is for on-time development of diagnostics, monitoring and therapeutic tools against virus in the global pandemics condition. Toward this goal, we have generated 18 high-quality whole-genome sequence data from 32 SARS-CoV-2 genotypes of PCR-positive COVID-19 patients, sampled from the Tashkent region of Uzbekistan. The nucleotide polymorphisms in the sequenced sample genomes were determined, including nonsynonymous (missense) and synonymous mutations in coding regions of coronavirus genome. Phylogenetic analysis grouped fourteen whole genome sample sequences (1, 2, 4, 5, 8, 10-15, 17, 32) into the G clade (or GR sub-clade) and four whole genome sample sequences (3, 6, 25, 27) into the S clade. A total of 128 mutations were identified, consisting of 45 shared and 83 unique mutations. Collectively, nucleotide changes represented one unique frameshift mutation, four upstream region mutations, six downstream region mutations, 50 synonymous mutations, and 67 missense mutations. The sequence data, presented herein, is the first coronavirus genomic sequence data from the Republic of Uzbekistan, which should contribute to enrich the global coronavirus sequence database, helping in future comparative studies. More importantly, the sequenced genomic data of coronavirus genotypes of this study should be useful for comparisons, diagnostics, monitoring, and therapeutics of COVID-19 disease in local and regional levels.

## Introduction

Severe acute respiratory syndrome coronavirus 2 (SARS-CoV-2) has high human-to-human transmission capacity and is classified within the *Coronaviridae* family, specifically the *Betacoronavirus* genus. It includes severe acute respiratory syndrome coronavirus (SARS-CoV, identified in 2002) [1] and Middle East respiratory syndrome coronavirus (MERS-CoV, identified in 2012) [2]. Comparative analysis has shown that the SARS-CoV-2 spike protein has more than 10-fold higher binding activity compared to SARS-CoV [3]. These viruses infect bats and are transmitted to humans via zoonotic transmission [4].

The first cases of coronavirus disease 2019 (COVID-19) were identified in Wuhan, Hubei Province, China, in December 2019, after which the infectious disease has become distributed globally. Currently, variants of SARS-CoV-2, the causative agent of COVID-19, continue to proliferate rapidly. Although China attempted to reduce the distribution of SARS-CoV-2 within the country, the infection quickly was spread in parts of Hubei Province neighboring Wuhan [5].

Later, SARS-CoV-2 has become distributed worldwide, representing a serious global issue; in particular, Europe and Latin America have suffered more than other countries in the beginning of coronavirus pandemics. The World Health Organization (WHO) officially declared a pandemic on March 11, 2020. Overall, 161,513,458 confirmed cases of COVID-19 with 3,209,109 deaths, including 92,320 confirmed cases and 655 deaths in Uzbekistan, have been reported to the WHO as of May 4, 2021 [6].

Microscopic vision of SARS-CoV-2 is a spherical, enveloped particle, positive-sense, single stranded RNA. Its genome contains 29,903 nucleotides [7]. The SARS-CoV-2 genome contains the open reading frame (ORF) proteins, spike (S), envelope (E), membrane (M), and nucleocapsid (N) genes in a 5’-3’ orientation. The replication ORF1ab gene of SARS-CoV-2 is the longest among other genes, 21,291 nt in length, and contains 16 predicted non-structural proteins, followed by (at least) 13 downstream ORFs. Moreover, it shares a highly conserved domain (amino acids 122–130) in nsp1 with SARS-CoV-2. The other genes, such as the S, ORF3a, E, M, and N genes of SARS-CoV-2, are 3,822, 828, 228, 669, and 1,260 nt in length, respectively [8]. The SARS-CoV-2 S protein binds to the host receptor, enabling entry into the cell. The gene encoding this protein is an important part of the viral genome, along with high sequence variability as a key point for new mutations [7].

Full and partial genome sequences of SARS-CoV-2 obtained from different countries and laboratories are now available at the Global Initiative on Sharing All Influenza Data (GISAID) [9], the National Center for Biotechnology Information (NCBI) [10] and the Virus Pathogen Database and Analysis Resource (ViPR) [11]. Notably, SARS-CoV-2 variants have been studied by analyzing 48,635 complete SARS-CoV-2 genomes obtained from GISAID [12]. Additionally, SARS-CoV-2 mutations were annotated by comparison with the sequence of the Wuhan-Hu-1 SARS-CoV-2 isolate, a reference genome NC_045512.2, and an average of seven mutations per sample was detected. The analysis has shown that major mutational types worldwide are caused by single-nucleotide transitions [12].

To date, at least three clades have been characterized based on geographic and genomic specificity of SARS-CoV-2 variants. The first is the V clade, defined in the GISAID EpiCoV portal, with limited variability. This clade members were isolated in Asia, and their genomes are similar to the reference coronavirus genome NC_045512.2 [13]. The second is the G clade (branching into its two offspring subclades, GH and GR), whose members are scattered across Europe and on the East Coast of the USA. This clade members carry a D614G amino acid mutation in the S protein and four characteristic point mutations: C241T, C3037T, C14408T, and A23403G. The third is the S clade, whose members have mostly been isolated in the USA, especially on the West Coast, and is characterized by C3037T and T28144C mutations [14].

The global concern related to novel virus variants is based on their rapid mutation rates, helping viral evasion of immunity formed by approved vaccines or previous variant(s) and cause reinfection [15]. Among all mutations in the SARS-CoV-2 genome, S gene mutations are important because the spike protein defines the viral host range and is often the target of neutralizing antibodies [16,17]. For example, the D614G substitution in the S protein occurred due to the A23403G missense mutation resulting in an amino acid change from aspartate to a glycine residue at position 614 (D614G). This mutation has emerged as a predominant G clade in Europe (954 of 1,449 (66%) sequences).

Since the beginning of the pandemics, the coronavirus variant carrying the G614 mutation has been rapidly distributed across the world and has become the dominant global strain [18,19]. Even in local epidemics, the shift has occurred by converting the D614 to G614 variant. These changes were highly statistically significant, and the G614 variant may have a fitness and selective advantage [18]. On the other hand, if the mutation occurred early in the growing population, a genetic drift could also help to increase the frequency of a specific variant without selective advantage [20].

Moreover, other mutations (N439K in RBD) were observed in the S protein [21]. The major frequent mutations, such as D614G, N439K, and S477N, in the S protein are causing the increased viral transmissibility. A variant containing D614G-associated mutations in the RBD became more infective and increased resistance to some neutralizing antibodies, with obvious implications for the recovery of COVID-19 patients [22].

SARS-CoV-2 cases were first identified on March 15, 2020, in passengers returning from Europe to the capital city Tashkent, Republic of Uzbekistan. The government of Uzbekistan has begun a massive testing of people who had symptoms of COVID-19 infectious disease. This event was the beginning of the epidemic in Uzbekistan that required effective diagnostics and monitoring of rapidly spreading coronavirus genotypes among population.

Therefore, with the main goal of identifying virus genotypes distributed in our territory as well as studying genomic diversity, types of mutations and possible emergence of new variants of SARS-CoV-2, we initiated the whole genome sequencing of COVID-19 samples. Here, we present the first whole-genome sequence data from COVID-19-infected patients in the Republic of Uzbekistan. We successfully assembled the 18 high-quality sample genome sequences for coronavirus genotypes and profiled 128 mutations with nonsynonymous and synonymous types. Comparative analysis using globally known genomes of SARS-CoV-2 grouped Uzbekistan sample genomes into two major clades of S and GR in the global phylogenetic tree. The sequenced genomic data of coronavirus genotypes, described herein, should be useful in fighting against coronavirus threats in Uzbekistan.

## Materials and methods

### Sample collection

In the middle of October and beginning of December of 2020, samples were collected several times from one hundred symptomatic patients with high temperature and occasional cough using nasopharyngeal and oropharyngeal swabs sticks (Huachenyang Technology, Shenzhen, China) and immediately placed in viral transport medium. Patients with possible COIVID-19 infection were from the diagnostics laboratories of the Tashkent Region Epidemiological Centre and the private BiogenMed COVID-19 testing center, Tashkent, Republic of Uzbekistan.

Biological samples were collected randomly from PCR-positive patients after laboratory testing for SARS-CoV-2. The research study has been approved by the Ethics Committee under the Ministry of Health of the Republic of Uzbekistan (#6/20-1582). All the experiments were carried out in accordance with the relevant guidelines and regulations. Samples were renumbered and de-identified so no one, even researchers could know the identity of the patients. For the reporting purpose only anonymous data including age and biological sex were kept.

We received a verbal consent for a voluntary participation from all patients involved for sample collection. Fall participants we have explained the use of collected samples for a sequencing experiment only without disclosing their personal identity or disturbing them in future. Verbal consent was preferred to written consent in this study because patients were not in a mood of signing any written document because of worriedness about COVID-19 infection at that time and had a hesitation because of not full-understanding of genome sequencing experiment. No minors were involved for the sample collection. Because of non-invasiveness of sequencing experiment, non-involvement of participants to any further downstream clinical procedures as well as anonymity of the personal identity of samples in this study, there was no requirement to obtain a consent document approval by the Ethics Committee.

Total RNA was extracted from the collected samples using a MagMax Viral/Pathogen kit (Life technologies Corporation, Austin, Texas, USA). Each sample was tested by PCR systems (threshold – 0,050) to the presence of SARS-CoV-2 using a CFX Connect Real-Time PCR System (BioRad, Hercules, California, USA). RT-PCR were implemented in 40 μl of final volumes. PCR amplification was carried out by the following steps: for cDNA synthesis at 35°C for 20 min, a first denaturation at 95°C for 5 min followed by 50 cycles of 94°C for 15 sec and 64°C for 20 sec (set fluorescence measurement for Fam, Hex, Rox, and Cy5 channels at 64°С) and a SARS-CoV-2/SARS CoV multiplex real-time PCR assay targeting the nucleocapsid (N), envelope (E) and region of SARS-CoV-like viruses (DNA Technology, Moscow, Russia). Among all tested patients, 32 PCR-positive samples (18 females and 14 males) were selected for further studies, which were randomly selected.

### SARS-CoV-2 sequencing

Complementary DNA (cDNA) was synthesized from 5 μl of RNA sample using a SuperScript VILO with DNAse cDNA Synthesis Kit (Life Technologies, Carlsbad, California, USA) and a ProFlex™ Base (Life Technologies holding Pte Ltd, Mapletree, Singapore). Libraries were constructed manually using the Ion AmpliSeq SARS-CoV-2 Research Panels, Ion Xpress Barcodes, and an Ion AmpliSeq Library Kit Plus Life Technologies Corporation, Frederick, Maryland, USA) following the manufacturer’s recommendations; the process included using amplification cycles from 12- to 24,000-plex in a single well based on viral load. Template amplification and enrichment as part of the manual workflow for the Ion S5 systems were performed with the Ion OneTouch 2 system using an Ion 540 Kit (Life Technologies, Carlsbad, California, USA). The thirty-two samples were multiplexed on an Ion 540 chip and sequenced using an Ion GeneStudio S5 Semiconductor Sequencer (Life Technologies Holdings Pte Ltd., Singapore).

### Sequenced data analysis

Sequenced reads were aligned with the Wuhan-Hu-1 Reference Genome (NC_045512.2) on the Torrent Suite v. 5.12.2 (Life Technologies, Carlsbad, California, USA). Plugins were used as follows by order coverage analysis (v5.12.0.0) and Variant Caller (v.5.12.0.4), both with default parameters and COVID19AnnotateSnpED (v1.3.0.2; Life Technologies, Carlsbad, California, USA). To predict the effect of a base substitution, a plugin specifically developed for SARS-CoV-2. VCF files generated by Torrent Suite (Life Technologies, Carlsbad, California, USA). Variant caller was filtered to remove variants with read depths less than 1000 and ion torrent quality scores less than 400 to keep reliable variants only (S1 Table). The filtered variants were used for sample clustering with Maximum Likelihood Tree in Molecular Evolutionary Genetics Analysis (MEGA, https://www.megasoftware.net) software. The consensus for each SARS-CoV-2 genome sequence were then submitted to the GISAID [9] under the accession numbers of EPI_ISL_1402423 to EPI_ISL_1477049 (available for registered users) and NCBI under the accession numbers GI:2021275696 to GI:2021275839 (or MW853559.1 to MW853569.1) [10] databases (Table 1).

**Table 1.**
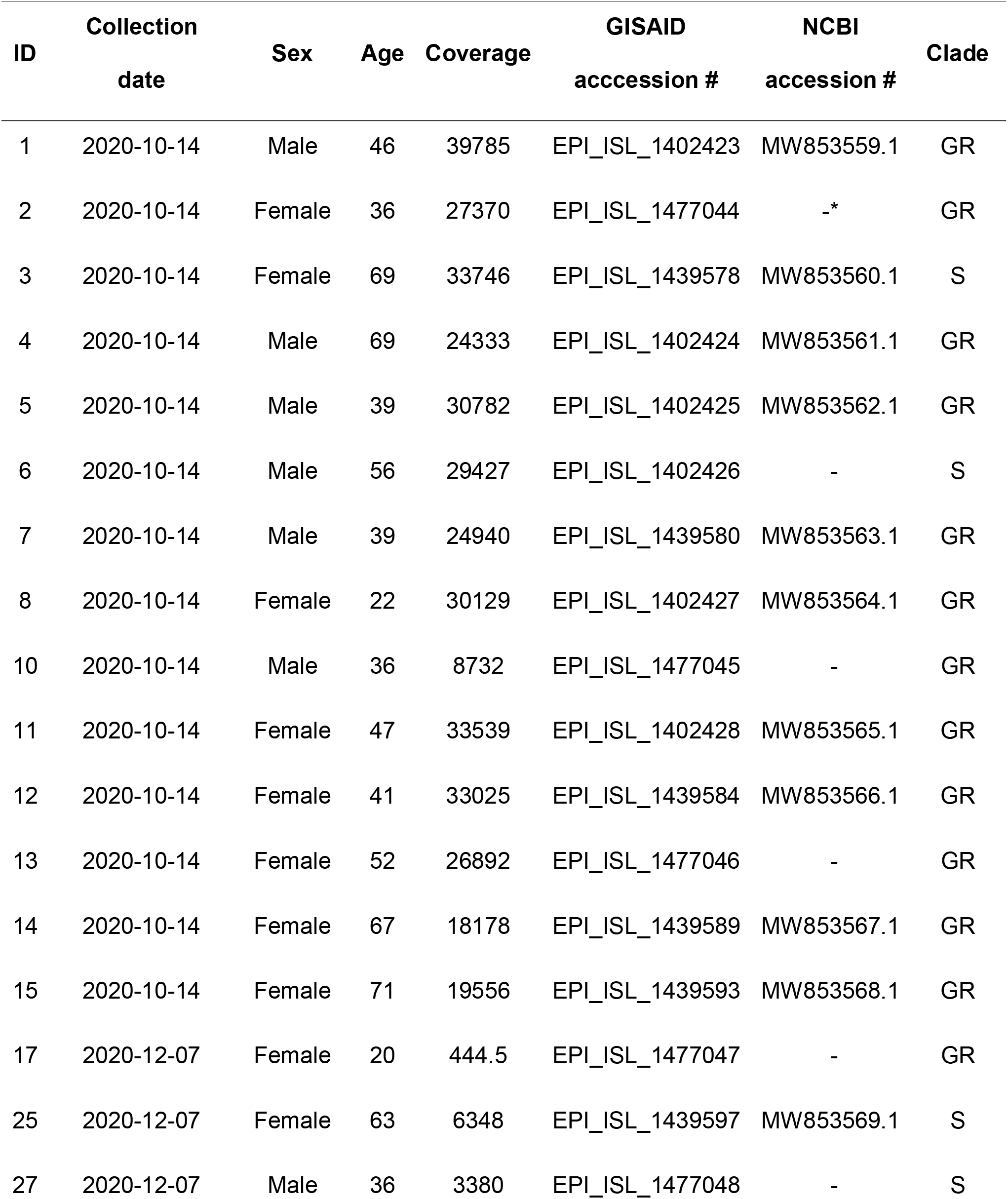
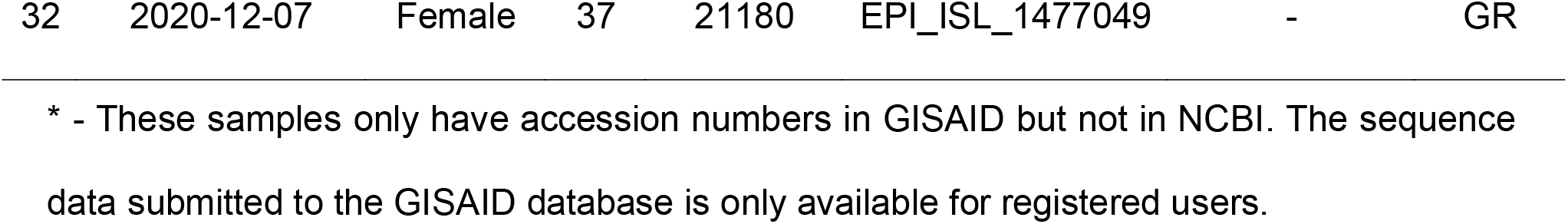
Summary data for the COVID-19 samples used in this study.

## Results

### Sample selection for sequencing

One hundred symptomatic patients with high temperature and occasional cough were collected for this study at the Tashkent Region Epidemiological Centre and a private COVID-19 testing center. Among them, 32 PCR-positive samples were selected for sequencing. There were seven men and eleven women with an average age of 47 (Table 1).

Out of 32 SARS-CoV-2 samples sequenced, 14 samples were excluded from further analysis due to the low quality of sequencing coverage, which resulted in several gaps in the consensus sequence. The remaining 18 samples (seven men and eleven women, Table 1) were selected with an average of mapped reads per sample of 3.9 million, with an average mean read depth of 23,203. However, the average uniformity of coverage in selected samples was low (75.31%; S2 Table).

### Analyzing the most reliable mutations among all sequenced samples

The analyzed mutations in these selected samples were generated by Variant Caller. According to the results, a number of mutations (Fig 1 and Fig 2) varied from five (samples 1,2 and 5) to 20 (samples 25 and 37). Most viral genomes contained between 8 and 15 mutations when compared with the NC_045512.2 reference genome, with mutations in sample 11 (15 mutations), sample 13 (14 mutations) and sample 10 and 15 (11 mutations), sample 8 (8 mutations) and sample 14 (9 mutations).

**Fig 1.**
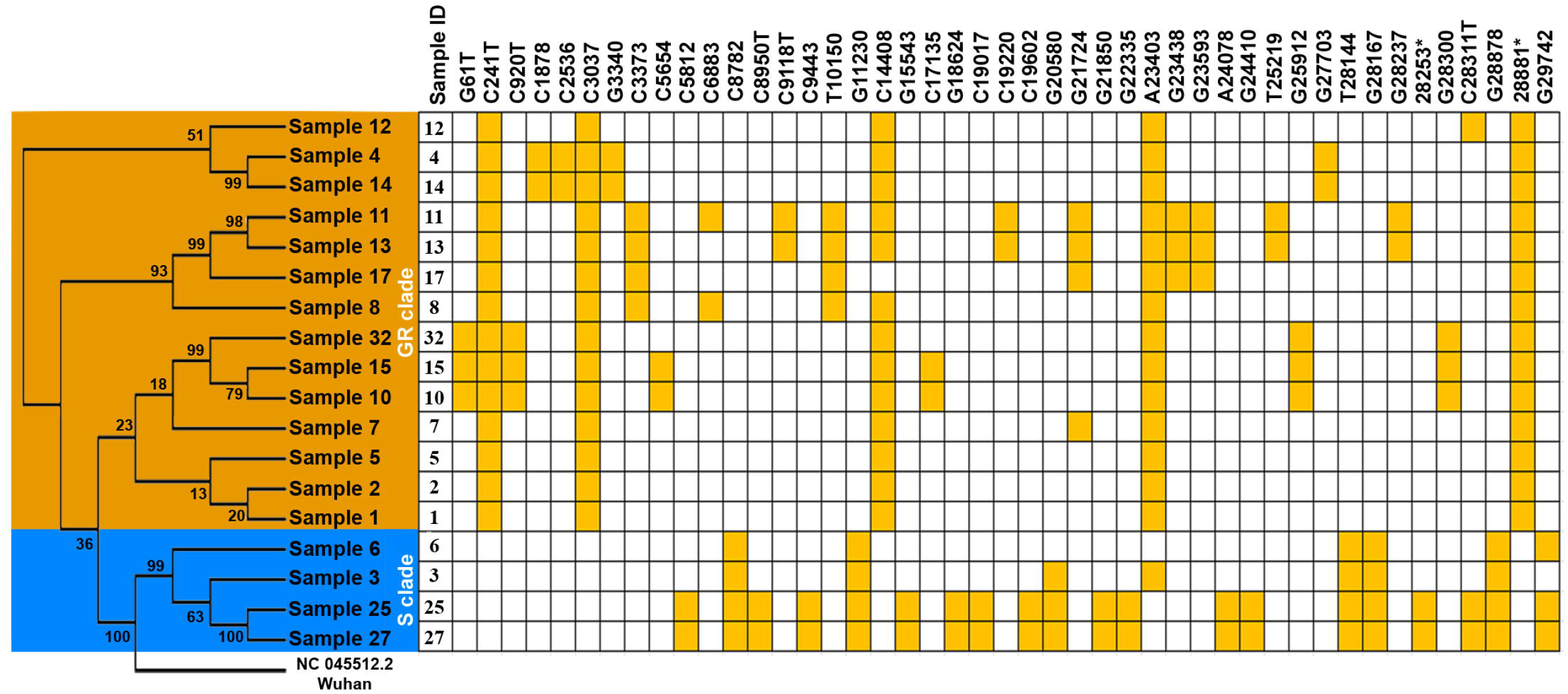
The Uzbek SARS-CoV-2 sequences clustered into 2 clades. A phylogenetic tree was generated in MEGA X based on the 18 viral sequences of severe acute respiratory syndrome coronavirus 2 (SARS-CoV-2) in samples collected from COVID-19 patients in the Tashkent region using the maximum likelihood method. This maximum likelihood tree was rooted by the Wuhan reference strain NC_045512.2 and produced using 128 mutations discovered in the 18 sequences (see S1 Table). The distribution of mutations shared by two or more sequences. A CA→TC substitution is located at position 28253*, and a GGG → AAC substitution is at 28881**. Bootstrap values are shown.

**Fig 2.**
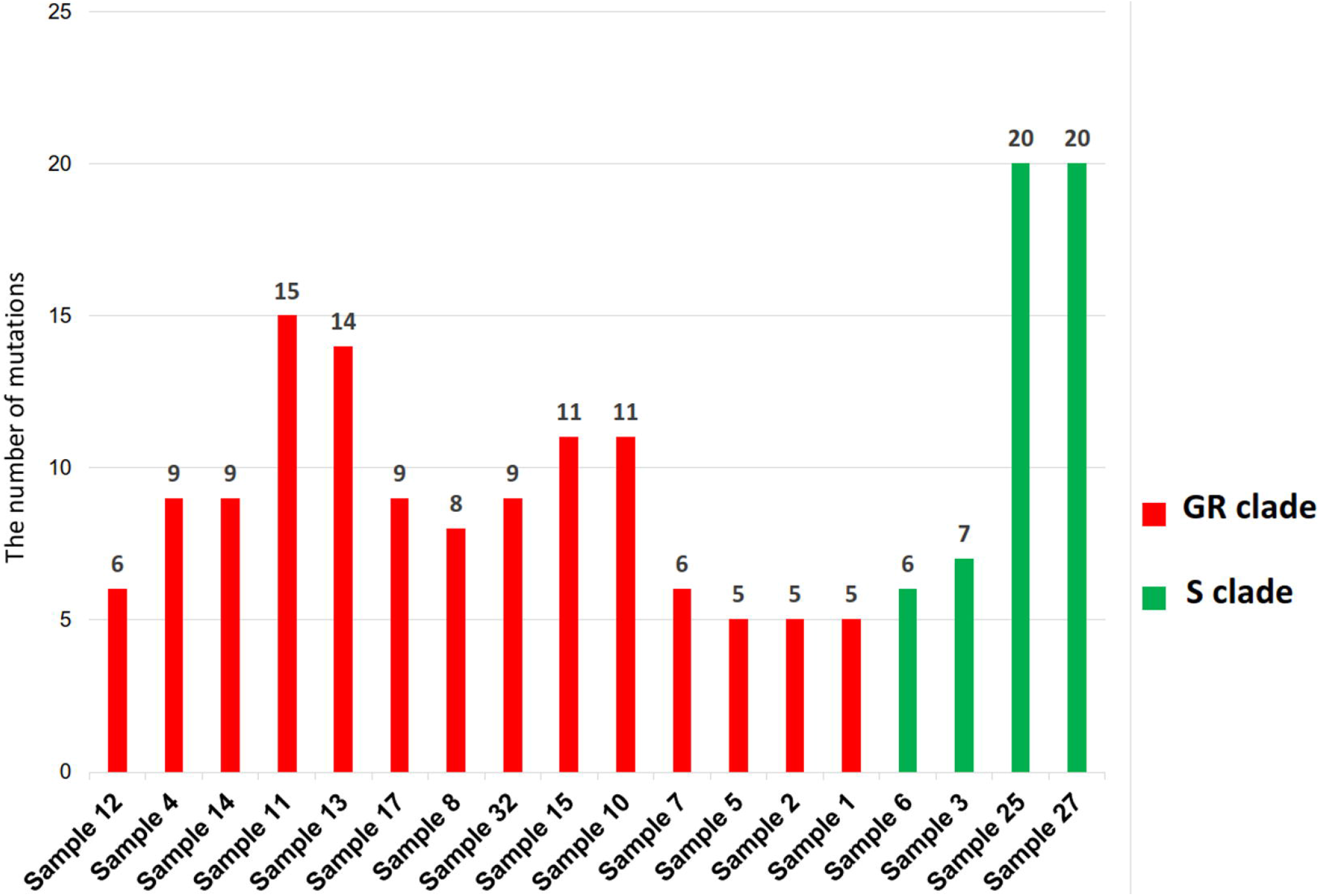
Number of mutations in COVID-19 samples sequenced. Note that the number of mutations is calculated per individual samples.

The most common nucleotide substitution detected (see S1 Table) was from cytosine to thymine (52/128 mutations), followed by guanine to thymine (36/128 mutations) and thymine to cytosine (10/128 mutations). All the shared mutations were homozygous. We observed one unique frameshift mutation (21574; c.13delC), seven shared mutations (G21850T, G22335T, A23403G, G23438T, G23593T, A24078G and G24410C), five unique (T22020C, T22478C, G22484T, C23634T and G24872T) missense mutations, two shared (G21724A and T25219G) and five unique (A23503T, C23758T, C24023T, G24199T and C24442T) synonymous mutations and five unique (A29676G, G29692T, C29708T and C29733T) with large deletions (4373delTCACCGAGGCCACGCGGAGTACGATCGAG) and one shared (G29742A) downstream region mutations in the gene encoding the S protein (S3 Tables). One missense mutation was identified in the E (envelope) region; two synonymous mutations were found in M (matrix), and one synonymous and eleven missense mutations were found in N (nucleocapsid).

Furthermore, 29 missense mutations, 36 synonymous mutations and three upstream gene mutations were found in the ORF1ab region. One synonymous mutation and seven missense mutations were detected in the ORF3a region. The ORF6 region showed one upstream mutation while the ORF7a region exhibited two missense and two synonymous mutations. Finally, five missense mutations and one synonymous mutation were found in the ORF8a region. Overall, we identified a total of 128 mutations, consisting of 45 shared and 83 unique mutations representing one unique frameshift mutation, four upstream region mutation, six downstream region mutation, 50 synonymous mutations, and 67 missense mutations (S1 Table).

### The phylogenetic tree was drawn based on viral sequences

We aimed to analyze the major mutations in all sequences to determine the difference between our cases and those worldwide. For this, a phylogenetic tree was generated in MEGA X based on the 18 viral sequences using the maximum likelihood method (Fig 1). One hundred twenty-eight mutations were obtained from eighteen SARS-CoV-2 viral genome sequences in samples from COVID-19 patients (S1 Table) by Variant caller (v.5.12.0.4).

The sequences have clustered into two large clades (S2 Table), including four sequences with five shared mutations that corresponded to the S clade (C8782T, G11230T, T28144C, G28167A and G28878A) and 14 samples with four mutations grouped in the GR subclade (C241T, C3037T, C14408T and A23403G). Two serial CA-TC substitutions (C28253T and A28254C) in some of the S clade viruses were identified (samples 25 and 27), and three serial GGG-AAC substitutions (G28881A, G28882A and G28883C) were identified in GR subclade viruses. In the GR subclade, two out-grouped samples (samples 2 and 5) and smaller clusters that corresponded to cluster 2 (samples 1 and 7) and cluster 3 (samples 4, 8, 10, 11, 12, 13, 14, 15, 17, and 32) were also found. In the S clade viruses, we observed one small cluster that corresponded to cluster 1 (samples 3, 6, 25 and 27).

Global phylogenetic tree was produced using local COVID-19 patient-derived sequences and the reference genome NC_045512.2, using nextstrain.org website (Fig 3). According to this analysis, the viruses in samples 3, 6, 25 and 27 belong to the S clade of SARS-CoV-2 and have sequences very similar to that of the reference genome, even though they harbor more substitutions in each gene, particularly in the S region. These variants also have grouped with African and Near East variants in the global phylogenetic tree provided by nextstrain.org. The remaining 14 sequences belong to the G clade, which possibly originated in Europe and North America [23].

**Fig 3.**
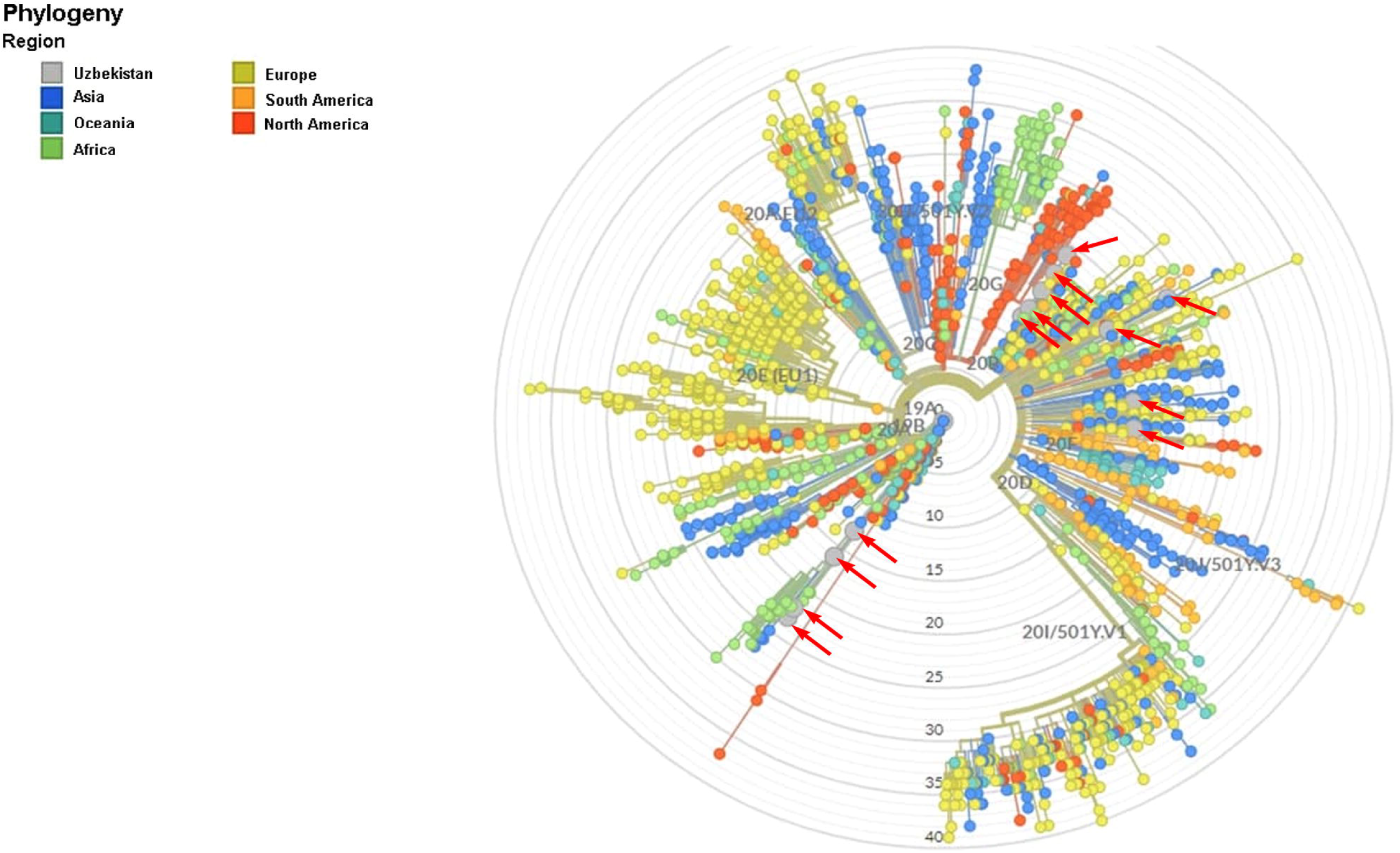
Global phylogenetic tree provided by nextstrain.org including COVID-19 samples from Uzbekistan. *Uzbekistan samples are shown with arrows, see S2 Table for variant and origin of Uzbekistan samples.

## Discussion

Our results revealed that whole-genome sequences of isolates obtained from the 18 symptomatic COVID-19 patients represent important nucleotide diversity (Table 1, S1-S3 Tables). These 18 sequenced samples have grouped in two major clades of SARS-CoV-2 on the public database of the GISAID named clade G (or GR subclade) and clade S according to the similarity of mutation signatures. The G clade, whose members are scattered across Europe and on the East Coast of the USA and carry a D614G amino acid mutation in the S protein and four characteristic point mutations namely C241T, C3037T, C14408T, and A23403G [12, 24, 25].

Among the 18 sequenced samples, fourteen samples have grouped into subclade GR. This branch contained an important deleterious trinucleotide mutation GGG28881AAC in the N gene of the coronavirus, inducing an ArgGly203LysArg change. This mutation later was found in many subsequent studies, with a frequency in deceased (39.45%) and recovered patients (31.38%) [25, 26], making them globally dominant in the coronavirus genomes. These 14 samples also contained well defined C28311T mutation of proline to leucine amino acid change in the N gene, playing a key role in the formation of replication– transcription complexes [25, 27] In addition, twenty shared mutations in these fourteen samples (G61T, C920T, C1878T, C2536T, G3340T, C3373A, C5654T, C6883T, C9118T, T10150C, C17135T, C19220T, G21724A, G23438T, G23593T, T25219G, G25912T, G27703A, G28237A, G28300T) were new and have not been described in any other studies before.

The remaining four samples out of 18 sequenced have grouped into the S clade and contained S clade-specific mutations such as C8782T and T28144C, including Leucine (L) to Serine (S) amino acid change (L84S) [28]. These mutations were found in many studies [12, 26, 28]. In addition, G11230T, G28167A, and G28878A mutations found in all four samples of our S clade were similar to Lui et al. [26], and were found in United Emirate Arabic. The G29742A mutation, shared by three samples out of four S clade samples (except Sample 3) was also found in United Emirate Arabic [26]. Other shared mutations found in four samples (C5812T, C8950T, C9443T, G15543T, G18624T, C19017T, C19602T, G20580T, G21850T, G22335T, A24078G, G24410C, 28253*, C28311T) were new and have not mentioned by any other studies. Moreover, we found a number of unique mutations in each sample that were specific for our SARS-CoV-2 genome sequence data.

Sequencing of the virus genome is fundamentally important for identifying SARS-CoV-2 strains and investigating local and global spread. In addition, the full-genome sequence of any virus that causes the infection can be helpful for investigating outbreak dynamics, such as changes in the size of the epidemic over time as well as spatiotemporal spread and transmission routes (WHO COVID report 2021). This is the first step attempt to sequence the full genome of SARS-CoV-2 from COVID-19-positive patients of the Republic of Uzbekistan. The first identified COVID-19 cases in Tashkent are believed to be of foreign origin due to international travel. Indeed, the results of this study showed that many of the infections originated from European (GR subclade) and Near East (S clade) countries. The cause of spreads was the result of international travelling.

The mutations between samples showed how virus structure changed itself over time when conditions became different. The genomic sequence data generated from these 18 samples, submitted to global databases such as NCBI and GISAID, should be helpful for public health and research organizations to observe the dynamics of disease spread in the region. Moreover, results will assist in the design of diagnostic assays, medications and vaccines. In this context, several institutions, including our centre, have already started working on a national vaccine based on these SARS-CoV-2 genome sequences found in symptomatic Uzbek patients.

Furthermore, the sequence data presented herein should add new sequence and mutational profile data from our region to a COVID sequence database (GISAID), useful for future molecular epidemiology and evolutionary phylogenetic studies by health and research organizations. In particular, our whole-genome sequence data, reported herein, should be helpful for tracking the origin and source of the currently spreading SARS-CoV-2 variants and for identifying and comparing the emerging new variants in Uzbekistan and beyond.

Here, we only provided the first effort of SARS-CoV-2 genome sequencing obtained from infected Uzbek patients using samples collected at the end of 2020. They were the early phase samples of the coronavirus disease pandemic that were spread to the whole country by that time. To date, the mutations multiplied by human-to-human transmission, and new strains have been scattered again in the country through international travels. The new mutations and variants require further sequencing efforts and analysis, which is in progress.

## Supporting information

**S1 Table. One hundred twenty-eight mutations were observed in eighteen SARS-CoV-2 viral genome sequences in samples from COVID-19 patients in Tashkent, Uzbekistan**.

**S2 Table. Genomes representing each of the major evolutionary lineages represented in our cohort**.

**S3 Table. Nucleotide mutations of the spike region of Uzbekistan SARS-CoV-2 sequences based on comparison to the reference sequence** (GenBank reference sequence accession number NC_045512.2).

## Acknowledgments

We thank the Sanitary-Epidemiological and Public Health Department of Tashkent Region, Ministry of Health of Uzbekistan, and the private clinic of BiogenMed, Tashkent, Uzbekistan, including but not limited to Mr. Abdukhakim M. Sotvoldiev, Mrs. Ra’no M. Abidova, Mrs. Larisa E. Alieva, Mr. Botir B. Sattorov and Ms. Saule A. Karimova, for their help in collection of the samples from symptomatic patients.

## Author Contributions

**Conceptualization &Data curation**: Mirzakamol S. Ayubov, Zabardast T. Buriev, Mukhammadjon K. Mirzakhmedov.

**Funding acquisition**: Mirzakamol S. Ayubov, Zabardast T. Buriev.

**Investigation, Methodology &Visualization**: Mirzakamol S. Ayubov, Mukhammadjon K. Mirzakhmedov, Abdurakhmon N. Yusupov, Dilshod E. Usmanov, Shukhrat E. Shermatov, Khurshida A. Ubaydullaeva.

**Supervision**: Ibrokhim Y. Abdurakhmonov, Zabardast T. Buriev.

**Writing – original draft**: Mirzakamol S. Ayubov, Mukhammadjon K. Mirzakhmedov, Zabardast T. Buriev.

**Writing – review & editing**: Ibrokhim Y. Abdurakhmonov, Mirzakamol S. Ayubov.

