## Supplementary Tables for "Profiling of the most reliable mutations from sequenced SARS-CoV-2 genomes scattered in Uzbekistan"

**S1 Table: One hundred twenty-eight mutations were observed in eighteen SARS-CoV-2 viral genome sequences in samples from COVID-19 patients in Tashkent, Uzbekistan.**

| **#** | **Mutation position** | **Genotype** | **Gene** | **HGVS_C*** | **HGVS_P**** | **MUTATION Effect** | **# of sequences carrying mutation** | **Ion Torrent quality score(-10logP)***** |
| --- | --- | --- | --- | --- | --- | --- | --- | --- |
| 1 | 61 | Homozygous | orf1ab | c.-205G>T | . | upstream region mutation | 3 | 2925.61 |
| 2 | 186 | Homozygous | orf1ab | c.-80C>T | . | upstream region mutation | 1 | 2944.42 |
| 3 | 241 | Homozygous | orf1ab | c.-25C>T | . | upstream region mutation | 14 | 2854.35 |
| 4 | 920 | Homozygous | orf1ab | c.655C>T | p.Leu219Leu | synonymous mutation | 3 | 2933.33 |
| 5 | 1594 | Homozygous | orf1ab | c.1329C>T | p.Ser443Ser | synonymous mutation | 1 | 2919.66 |
| 6 | 1648 | Homozygous | orf1ab | c.1383C>T | p.Ile461Ile | synonymous mutation | 1 | 2962.64 |
| 7 | 1878 | Homozygous | orf1ab | c.1613C>T | p.Ser538Leu | missense mutation | 2 | 2976.88 |
| 8 | 2437 | Homozygous | orf1ab | c.2172A>T | p.Arg724Ser | missense mutation | 1 | 2970.73 |
| 9 | 2536 | Homozygous | orf1ab | c.2271C>T | p.Val757Val | synonymous mutation | 2 | 2950.37 |
| 10 | 2886 | Homozygous | orf1ab | c.2621T>C | p.Val874Ala | missense mutation | 1 | 2919.66 |
| 11 | 3037 | Homozygous | orf1ab | c.2772C>T | p.Phe924Phe | synonymous mutation | 14 | 2950.45 |
| 12 | 3340 | Homozygous | orf1ab | c.3075G>T | p.Val1025Val | synonymous mutation | 2 | 2975.51 |
| 13 | 3373 | Homozygous | orf1ab | c.3108C>A | p.Asp1036Glu | missense mutation | 4 | 2910.5 |
| 14 | 4582 | Homozygous | orf1ab | c.4317C>T | p.Asn1439Asn | synonymous mutation | 1 | 2974.56 |
| 15 | 4898 | Homozygous | orf1ab | c.4633C>T | p.His1545Tyr | missense mutation | 1 | 2966.91 |
| 16 | 5128 | Homozygous | orf1ab | c.4863A>G | p.Leu1621Leu | synonymous mutation | 1 | 2974.23 |
| 17 | 5175 | Homozygous | orf1ab | c.4910C>T | p.Thr1637Ile | missense mutation | 1 | 2935.9 |
| 18 | 5221 | Homozygous | orf1ab | c.4956C>T | p.His1652His | synonymous mutation | 1 | 2976.87 |
| 19 | 5386 | Homozygous | orf1ab | c.5121T>C | p.Ala1707Ala | synonymous mutation | 1 | 2943.24 |
| 20 | 5654 | Homozygous | orf1ab | c.5389C>T | p.Leu1797Leu | synonymous mutation | 2 | 2881.29 |
| 21 | 5812 | Homozygous | orf1ab | c.5547C>T | p.Asp1849Asp | synonymous mutation | 2 | 2962.04 |
| 22 | 5826 | Homozygous | orf1ab | c.5561C>T | p.Thr1854Ile | missense mutation | 1 | 2968.67 |
| 23 | 5974 | Homozygous | orf1ab | c.5709C>T | p.Asp1903Asp | synonymous mutation | 1 | 1451.89 |
| 24 | 6883 | Homozygous | orf1ab | c.6618C>T | p.Val2206Val | synonymous mutation | 2 | 2171.77 |
| 25 | 8645 | Homozygous | orf1ab | c.8380C>T | p.His2794Tyr | missense mutation | 1 | 2934.35 |
| 26 | 8782 | Homozygous | orf1ab | c.8517C>T | p.Ser2839Ser | synonymous mutation | 4 | 2886.79 |
| 27 | 8950 | Homozygous | orf1ab | c.8685C>T | p.Ile2895Ile | synonymous mutation | 2 | 2911.18 |
| 28 | 9118 | Homozygous | orf1ab | c.8853C>T | p.Asp2951Asp | synonymous mutation | 2 | 2823.98 |
| 29 | 9443 | Homozygous | orf1ab | c.9178C>T | p.Leu3060Phe | missense mutation | 2 | 2627.79 |
| 30 | 10150 | Homozygous | orf1ab | c.9885T>C | p.Leu3295Leu | synonymous mutation | 4 | 2966.18 |
| 31 | 11144 | Homozygous | orf1ab | c.10879A>G | p.Met3627Val | missense mutation | 1 | 674.85 |
| 32 | 11230 | Homozygous | orf1ab | c.10965G>T | p.Met3655Ile | missense mutation | 4 | 2972.25 |
| 33 | 11298 | Homozygous | orf1ab | c.11033A>G | p.Lys3678Arg | missense mutation | 1 | 2412.26 |
| 34 | 11339 | Homozygous | orf1ab | c.11074C>T | p.Leu3692Leu | synonymous mutation | 1 | 2940.52 |
| 35 | 11417 | Homozygous | orf1ab | c.11152G>T | p.Val3718Phe | missense mutation | 1 | 2919.47 |
| 36 | 11446 | Homozygous | orf1ab | c.11181A>G | p.Leu3727Leu | synonymous mutation | 1 | 2937.18 |
| 37 | 11761 | Homozygous | orf1ab | c.11496G>T | p.Lys3832Asn | missense mutation | 1 | 2452.44 |
| 38 | 12148 | Homozygous | orf1ab | c.11883A>C | p.Gln3961His | missense mutation | 1 | 2921.54 |
| 39 | 12180 | Homozygous | orf1ab | c.11915A>G | p.Asp3972Gly | missense mutation | 1 | 2912.17 |
| 40 | 12191 | Homozygous | orf1ab | c.11926G>T | p.Val3976Phe | missense mutation | 1 | 2928.97 |
| 41 | 12202 | Homozygous | orf1ab | c.11937G>T | p.Lys3979Asn | missense mutation | 1 | 2907.64 |
| 42 | 12763 | Homozygous | orf1ab | c.12498C>T | p.Asp4166Asp | synonymous mutation | 1 | 2952.73 |
| 43 | 12784 | Homozygous | orf1ab | c.12519C>T | p.Asn4173Asn | synonymous mutation | 1 | 2830.87 |
| 44 | 13348 | Homozygous | orf1ab | c.13083G>T | p.Val4361Val | synonymous mutation | 1 | 2933.15 |
| 45 | 13365 | Homozygous | orf1ab | c.13100A>C | p.Asn4367Thr | missense mutation | 1 | 2437.84 |
| 46 | 13533 | Homozygous | orf1ab | c.13269A>G | p.Val4423Val | synonymous mutation | 1 | 2944.02 |
| 47 | 14371 | Homozygous | orf1ab | c.14107G>T | p.Ala4703Ser | missense mutation | 1 | 2977.02 |
| 48 | 14408 | Homozygous | orf1ab | c.14144C>T | p.Pro4715Leu | missense mutation | 13 | 2835.86 |
| 49 | 15048 | Homozygous | orf1ab | c.14784C>T | p.Ile4928Ile | synonymous mutation | 1 | 2339.25 |
| 50 | 15543 | Homozygous | orf1ab | c.15279G>T | p.Thr5093Thr | synonymous mutation | 2 | 2980.13 |
| 51 | 15546 | Homozygous | orf1ab | c.15282C>T | p.Ala5094Ala | synonymous mutation | 1 | 2928.36 |
| 52 | 16338 | Homozygous | orf1ab | c.16074C>T | p.Val5358Val | synonymous mutation | 1 | 2982.14 |
| 53 | 16943 | Homozygous | orf1ab | c.16679G>T | p.Ser5560Ile | missense mutation | 1 | 2942.39 |
| 54 | 17004 | Homozygous | orf1ab | c.16740C>T | p.Leu5580Leu | synonymous mutation | 1 | 2028.19 |
| 55 | 17135 | Homozygous | orf1ab | c.16871C>T | p.Pro5624Leu | missense mutation | 2 | 2818.32 |
| 56 | 18624 | Homozygous | orf1ab | c.18360G>T | p.Met6120Ile | missense mutation | 2 | 2981.39 |
| 57 | 19017 | Homozygous | orf1ab | c.18753C>T | p.Phe6251Phe | synonymous mutation | 2 | 2428.65 |
| 58 | 19220 | Homozygous | orf1ab | c.18956C>T | p.Ala6319Val | missense mutation | 2 | 2944.37 |
| 59 | 19497 | Homozygous | orf1ab | c.19233T>C | p.His6411His | synonymous mutation | 1 | 2948.86 |
| 60 | 19549 | Homozygous | orf1ab | c.19285G>T | p.Ala6429Ser | missense mutation | 1 | 2982.14 |
| 61 | 19602 | Homozygous | orf1ab | c.19338C>T | p.Asn6446Asn | synonymous mutation | 2 | 2980.27 |
| 62 | 19735 | Homozygous | orf1ab | c.19471G>T | p.Asp6491Tyr | missense mutation | 1 | 2976.29 |
| 63 | 19813 | Homozygous | orf1ab | c.19549C>T | p.Pro6517Ser | missense mutation | 1 | 2751.35 |
| 67 | 19839 | Homozygous | orf1ab | c.19575T>C | p.Asn6525Asn | synonymous mutation | 1 | 2724.84 |
| 65 | 20580 | Homozygous | orf1ab | c.20316G>T | p.Val6772Val | synonymous mutation | 3 | 2954.88 |
| 66 | 21255 | Homozygous | orf1ab | c.20991G>C | p.Ala6997Ala | synonymous mutation | 1 | 2919.18 |
| 67 | 21471 | Homozygous | orf1ab | c.21207T>C | p.Ile7069Ile | synonymous mutation | 1 | 2055.02 |
| 68 | 21520 | Homozygous | orf1ab | c.21256G>A | p.Val7086Ile | missense mutation | 1 | 2981.03 |
| 69 | 21574 | Homozygous | S | c.13delC | p.Val6fs | frameshift mutation | 1 | 2885.88 |
| 70 | 21724 | Homozygous | S | c.162G>A | p.Leu54Leu | synonymous mutation | 4 | 2950.01 |
| 71 | 21850 | Homozygous | S | c.288G>T | p.Glu96Asp | missense mutation | 2 | 2977.75 |
| 72 | 22020 | Homozygous | S | c.458T>C | p.Met153Thr | missense mutation | 1 | 2838.4 |
| 73 | 22335 | Homozygous | S | c.773G>T | p.Trp258Leu | missense mutation | 2 | 2882.61 |
| 74 | 22478 | Homozygous | S | c.916T>C | p.Phe306Leu | missense mutation | 1 | 2671.18 |
| 75 | 22484 | Homozygous | S | c.922G>T | p.Val308Leu | missense mutation | 1 | 2981.34 |
| 76 | 23403 | Homozygous | S | c.1841A>G | p.Asp614Gly | missense mutation | 15 | 2958.72 |
| 77 | 23438 | Homozygous | S | c.1876G>T | p.Ala626Ser | missense mutation | 3 | 2978.37 |
| 78 | 23503 | Homozygous | S | c.1941A>T | p.Ala647Ala | synonymous mutation | 1 | 2929.27 |
| 79 | 23593 | Homozygous | S | c.2031G>T | p.Gln677His | missense mutation | 3 | 2905.65 |
| 80 | 23634 | Homozygous | S | c.2072C>T | p.Ser691Phe | missense mutation | 1 | 2962.83 |
| 81 | 23758 | Homozygous | S | c.2196C>T | p.Thr732Thr | synonymous mutation | 1 | 2388.57 |
| 82 | 24023 | Homozygous | S | c.2461C>T | p.Leu821Leu | synonymous mutation | 1 | 2918.63 |
| 83 | 24078 | Homozygous | S | c.2516A>G | p.Asp839Gly | missense mutation | 2 | 2939.33 |
| 84 | 24199 | Homozygous | S | c.2637G>T | p.Ala879Ala | synonymous mutation | 1 | 2894.64 |
| 85 | 24410 | Homozygous | S | c.2848G>C | p.Asp950His | missense mutation | 2 | 2942.52 |
| 86 | 24442 | Homozygous | S | c.2880C>T | p.Asn960Asn | synonymous mutation | 1 | 2982.19 |
| 87 | 24872 | Homozygous | S | c.3310G>T | p.Val1104Leu | missense mutation | 1 | 2890.8 |
| 88 | 25219 | Homozygous | S | c.3657T>G | p.Gly1219Gly | synonymous mutation | 2 | 2832.51 |
| 89 | 25513 | Homozygous | ORF3a | c.121C>T | p.Leu41Phe | missense mutation | 1 | 2513.33 |
| 90 | 25613 | Homozygous | ORF3a | c.221C>T | p.Ser74Phe | missense mutation | 1 | 2908.51 |
| 91 | 25690 | Homozygous | ORF3a | c.298G>T | p.Gly100Cys | missense mutation | 1 | 587 |
| 92 | 25906 | Homozygous | ORF3a | c.514G>T | p.Gly172Cys | missense mutation | 1 | 2980.12 |
| 93 | 25912 | Homozygous | ORF3a | c.520G>T | p.Gly174Cys | missense mutation | 3 | 2967.31 |
| 94 | 26028 | Homozygous | ORF3a | c.636C>T | p.Tyr212Tyr | synonymous mutation | 1 | 2941.24 |
| 95 | 26135 | Homozygous | ORF3a | c.743C>T | p.Thr248Ile | missense mutation | 1 | 2951.24 |
| 96 | 26185 | Homozygous | ORF3a | c.793G>T | p.Asp265Tyr | missense mutation | 1 | 2978.71 |
| 97 | 26428 | Homozygous | E | c.184G>T | p.Val62Phe | missense mutation | 1 | 2930.42 |
| 98 | 26681 | Homozygous | M | c.159C>T | p.Phe53Phe | synonymous mutation | 1 | 2858.16 |
| 99 | 26951 | Homozygous | M | c.429G>T | p.Val143Val | synonymous mutation | 1 | 2905.14 |
| 100 | 27198 | Homozygous | ORF6 | c.-4A>T | . | upstream region mutation | 1 | 2893.69 |
| 101 | 27513 | Homozygous | ORF7a | c.120C>T | p.Tyr40Tyr | synonymous mutation | 1 | 2948.94 |
| 102 | 27682 | Homozygous | ORF7a | c.289T>C | p.Tyr97His | missense mutation | 1 | 2940.86 |
| 103 | 27684 | Homozygous | ORF7a | c.291C>T | p.Tyr97Tyr | synonymous mutation | 1 | 2972.71 |
| 104 | 27703 | Homozygous | ORF7a | c.310G>A | p.Val104Ile | missense mutation | 3 | 2365.53 |
| 105 | 28001 | Homozygous | ORF8 | c.108G>T | p.Pro36Pro | synonymous mutation | 1 | 2980.68 |
| 106 | 28144 | Homozygous | ORF8 | c.251T>C | p.Leu84Ser | missense mutation | 4 | 2980.56 |
| 107 | 28167 | Homozygous | ORF8 | c.274G>A | p.Glu92Lys | missense mutation | 4 | 1771.62 |
| 108 | 28178 | Homozygous | ORF8 | c.285G>T | p.Leu95Phe | missense mutation | 1 | 2746 |
| 109 | 28237 | Homozygous | ORF8 | c.344G>A | p.Arg115His | missense mutation | 2 | 2933.08 |
| 110 | 28253 | Homozygous | ORF8 | c.360_361delCAinsTC | p.Ile121Leu | missense mutation | 2 | 2893.15 |
| 111 | 28300 | Homozygous | N | c.27G>T | p.Gln9His | missense mutation | 3 | 2971.69 |
| 112 | 28311 | Homozygous | N | c.38C>T | p.Pro13Leu | missense mutation | 3 | 2907.61 |
| 113 | 28371 | Homozygous | N | c.98G>T | p.Ser33Ile | missense mutation | 1 | 2941.09 |
| 114 | 28378 | Homozygous | N | c.105G>T | p.Ala35Ala | synonymous mutation | 2 | 2900.07 |
| 115 | 28677 | Homozygous | N | c.404C>T | p.Thr135Ile | missense mutation | 1 | 2938.85 |
| 116 | 28851 | Homozygous | N | c.578G>T | p.Ser193Ile | missense mutation | 1 | 2921.23 |
| 117 | 28878 | Homozygous | N | c.605G>A | p.Ser202Asn | missense mutation | 4 | 2970.24 |
| 118 | 28881 | Homozygous | N | c.608_610delGGGinsAAC | p.ArgGly203LysArg | missense mutation | 14 | 2897.34 |
| 119 | 28887 | Homozygous | N | c.614C>T | p.Thr205Ile | missense mutation | 1 | 2925.39 |
| 120 | 28905 | Homozygous | N | c.632C>T | p.Ala211Val | missense mutation | 1 | 2919.09 |
| 121 | 28975 | Homozygous | N | c.702G>C | p.Met234Ile | missense mutation | 1 | 2912.27 |
| 122 | 29195 | Homozygous | N | c.922G>T | p.Ala308Ser | missense mutation | 1 | 2949.45 |
| 123 | 29676 | Homozygous | S | c.*4292A>G | . | downstream region mutation | 1 | 2682.06 |
| 124 | 29692 | Homozygous | S | c.*4308G>T | . | downstream region mutation | 1 | 2911.53 |
| 125 | 29708 | Homozygous | S | c.*4324C>T | . | downstream region mutation | 1 | 2892.98 |
| 126 | 29728 | Homozygous | S | c.*4345_*4373delTCACCGAGGCCACGCGGAGTACGATCGAG | . | downstream region mutation | 1 | 2907.15 |
| 127 | 29733 | Homozygous | S | c.*4349C>T | . | downstream region mutation | 1 | 2751.22 |
| 128 | 29742 | Homozygous | S | c.*4358G>A | . | downstream region mutation | 3 | 2955.28 |

***HGVS_C – Human Genome Variation Society, C*- coding DNA;**

****HGVS_P - Human Genome Variation Society, P**-protein sequence;**

*****Ion Torrent quality score (-10logP) – obtained by Variant Caller.**

**S2 Table: Genomes representing each of the major evolutionary lineages represented in our cohort.**

| **#** | **Sample ID** | **GISAID ID** | **Virus name** | **Collection date** | **Lineage (GISAID Clade)** | **Originating laboratory** |
| --- | --- | --- | --- | --- | --- | --- |
| **1** | **1** | EPI_ISL_1402423 | hCoV-19/Uzbekistan/Tashkent-CGB-01/2021 | 2020-10-14 | B.1.1.317 (GR) | Private clinic of BiogenMed, Tashkent, Uzbekistan |
| **2** | **2** | EPI_ISL_1477044 | hCoV-19/Uzbekistan/Tashkent-CGB-02/2020 | 2020-10-14 | B.1.1 (GR) | Private clinic of BiogenMed, Tashkent, Uzbekistan |
| **3** | **3** | EPI_ISL_1439578 | hCoV-19/Uzbekistan/Tashkent-CGB-03/2021 | 2020-10-14 | A.24 (S) | Private clinic of BiogenMed, Tashkent, Uzbekistan |
| **4** | **4** | EPI_ISL_1402424 | hCoV-19/Uzbekistan/Tashkent-CGB-04/2021 | 2020-10-14 | B.1.1 (GR) | Private clinic of BiogenMed, Tashkent, Uzbekistan |
| **5** | **5** | EPI_ISL_1402425 | hCoV-19/Uzbekistan/Tashkent-CGB-05/2021 | 2020-10-14 | B.1.1 (GR) | Private clinic of BiogenMed, Tashkent, Uzbekistan |
| **6** | **6** | EPI_ISL_1402426 | hCoV-19/Uzbekistan/Tashkent-CGB-06/2021 | 2020-10-14 | A (S) | Private clinic of BiogenMed, Tashkent, Uzbekistan |
| **7** | **7** | EPI_ISL_1439580 | hCoV-19/Uzbekistan/Tashkent-CGB-07/2021 | 2020-10-14 | B.1.1 (GR) | Private clinic of BiogenMed, Tashkent, Uzbekistan |
| **8** | **8** | EPI_ISL_1402427 | hCoV-19/Uzbekistan/Tashkent-CGB-08/2021 | 2020-10-14 | B.1.1.294 (GR) | Private clinic of BiogenMed, Tashkent, Uzbekistan |
| **9** | **10** | EPI_ISL_1477045 | hCoV-19/Uzbekistan/Tashkent-CGB-10/2020 | 2020-10-14 | B.1.1.274 (GR) | Private clinic of BiogenMed, Tashkent, Uzbekistan |
| **10** | **11** | EPI_ISL_1402428 | hCoV-19/Uzbekistan/Tashkent-CGB-11/2021 | 2020-10-14 | B.1.1.294 (GR) | Private clinic of BiogenMed, Tashkent, Uzbekistan |
| **11** | **12** | EPI_ISL_1439584 | hCoV-19/Uzbekistan/Tashkent-CGB-12/2021 | 2020-10-14 | B.1.1 (GR) | Private clinic of BiogenMed, Tashkent, Uzbekistan |
| **12** | **13** | EPI_ISL_1477046 | hCoV-19/Uzbekistan/Tashkent-CGB-13/2020 | 2020-10-14 | B.1.1.294 (GR) | Private clinic of BiogenMed, Tashkent, Uzbekistan |
| **13** | **14** | EPI_ISL_1439589 | hCoV-19/Uzbekistan/Tashkent-CGB-14/2021 | 2020-10-14 | B.1.1 (GR) | Private clinic of BiogenMed, Tashkent, Uzbekistan |
| **14** | **15** | EPI_ISL_1439593 | hCoV-19/Uzbekistan/Tashkent-CGB-15/2021 | 2020-10-14 | B.1.1.274 (GR) | Private clinic of BiogenMed, Tashkent, Uzbekistan |
| **15** | **17** | EPI_ISL_1477047 | hCoV-19/Uzbekistan/Tashkent-CGB-17/2020 | 2020-12-07 | B.1.1.294 (GR) | Sanitary-Epidemiological and Public Health Department of Tashkent Region, Uzbekistan |
| **16** | **25** | EPI_ISL_1439597 | hCoV-19/Uzbekistan/Tashkent-CGB-25/2021 | 2020-12-07 | A.24 (S) | Sanitary-Epidemiological and Public Health Department of Tashkent Region, Uzbekistan |
| **17** | **27** | EPI_ISL_1477048 | hCoV-19/Uzbekistan/Tashkent-CGB-27/2020 | 2020-12-07 | A.24 (S) | Sanitary-Epidemiological and Public Health Department of Tashkent Region, Uzbekistan |
| **18** | **32** | EPI_ISL_1477049 | hCoV-19/Uzbekistan/Tashkent-CGB-32/2020 | 2020-12-07 | B.1.1.274 (GR) | Sanitary-Epidemiological and Public Health Department of Tashkent Region, Uzbekistan |

**S3 Table. Nucleotide mutations of the spike region of Uzbekistan SARS-CoV-2 sequences based on comparison to the reference sequence** (GenBank reference sequence accession number NC_045512.2).

| **#** | **Nucleotide position** | **Reference nucleotide** | **Sequenced nucleotide** | **Mutation type** | **Nucleotide change** | **Amino acid change (position)** |
| --- | --- | --- | --- | --- | --- | --- |
| 1 | 21850 | G | T | missense | G→T | E96D |
| 2 | 22335 | G | T | missense | G→T | W258 L |
| 3 | 24078 | A | G | missense | A→G | D839G |
| 4 | 24410 | G | C | missense | G→C | D950H |
| 5 | 23403 | A | G | missense | A→G | D614G |
| 6 | 23438 | G | T | missense | G→T | A626S |
| 7 | 23593 | G | T | missense | G→T | Q677H |
| 8 | 22020 | T | C | missense | T→C | M153T |
| 9 | 24872 | G | T | missense | G→T | V1104 L |
| 10 | 22478 | T | C | missense | T→C | F306 L |
| 11 | 22484 | G | T | missense | G→T | V308 L |
| 12 | 23634 | C | T | missense | C→T | S691F |
| 13 | 21724 | G | A | synonymous | G→A | L54 L |
| 14 | 25219 | T | G | synonymous | T→G | G1219G |
| 15 | 23503 | A | T | synonymous | A→T | A647A |
| 16 | 23758 | C | T | synonymous | C→T | T732T |
| 17 | 24023 | C | T | synonymous | C→T | L821 L |
| 18 | 24199 | G | T | synonymous | G→T | A879A |
| 19 | 24442 | C | T | synonymous | C→T | N960N |
| 20 | 29676 | A | G | downstream region | A→G | - |
| 21 | 29692 | G | T | downstream region | G→T | - |
| 22 | 29708 | C | T | downstream region | C→T | - |
| 23 | 29733 | C | T | downstream region | C→T | - |
| 24 | 29742 | G | A | downstream region | G→A | - |
| 25 | 29728 | TCACCGAGGCCACGCGGAGTACGATCGAG | - | downstream region | *4345_*4373del | - |
